# Nanostructured Zirconia thin films as neurogliomorphic interface for neural cells of central and peripheral nervous system

**DOI:** 10.64898/2026.05.26.727630

**Authors:** Giorgia Conte, Francesca Borghi, Chiara Lazzarini, Claudio Piazzoni, Aikaterini Konstantoulaki, Roberta Fabbri, Marco Caprini, Paolo Milani, Valentina Benfenati

## Abstract

Recent advances in neuroscience have highlighted the central role of glial cells, particularly astrocytes, in regulating neural network activity through calcium-dependent neuron–glia communication. In parallel, nanostructured cluster-assembled materials have emerged as promising candidates for developing brain-machine interfaces, because of their biomimetic morphology, mechanotransductive properties and neuromorphic behavior. Among these, nanostructured zirconium oxide (ns-ZrOx) thin films have recently demonstrated memristive and signal-processing capabilities compatible with biohybrid neural systems, yet their interaction with heterogeneous neuroglial networks remains poorly understood.

Here, we investigate the biocompatibility and functional effects of ns-ZrOx interfaces on primary astrocytes and dorsal root ganglion (DRG) neuron–glia co-cultures, comparing nanostructured and flat zirconia substrates. Both substrates supported cellular adhesion, survival, and differentiation. However, ns-ZrOx selectively enhanced glial calcium signaling, increasing transient amplitude and accelerating response kinetics in both central and peripheral glial populations. Our findings identify ns-ZrOx as an active neurogliomorphic interface capable of modulating neuron–glia communication through nanoscale material properties. By bridging glial physiology with neuromorphic nanomaterials, this work supports the development of hybrid bioelectronic platforms integrating living neural networks with adaptive functional materials for brain-inspired computing and advanced neural interfaces.

## INTRODUCTION

Recent neuroscientific discoveries have profoundly reshaped our understanding of brain function, underscoring the crucial role of glial cells—particularly astrocytes—in the dynamic regulation of neural circuits^1,2^. Far from being passive support elements, astrocytes actively participate in synaptic transmission, modulate neuronal excitability, and maintain homeostasis through complex bidirectional signaling with neurons^3,4^. Their involvement in synaptic plasticity, metabolic regulation, and even information processing has positioned them as indispensable players in the orchestration of brain network behaviour. In particular, intracellular Calcium signaling ([Ca^2+^]_i_) of astrocytes is central to both neuronal excitability and astrocytic function, acting as a key messenger in synaptic transmission, gliotransmission, and network synchronization^2,5,6^. This paradigm shift highlights an urgent need for a new generation of brain-computer interface and devices, that counts on materials capable of interacting with heterogeneous neuronal elements, such as neurons but also with glial components.

Nanostructured cluster-assembled materials are good candidate to support and interact with heterogeneous neuronal culture, thanks to their mechanotrasductive^7–9^ and neuromorphic behaviours^10–12^. Mechanotransductive processes are promoted by the nanoscale structure of the cluster-assembled materials and its organization at the mesoscale^13^, with a morphology closely resembling the Extra Cellular Matrix (ECM) one^14,15^, thus providing biocompatible and bio-mimetic surfaces already shown to support neuronal and glial cultures^7,10^. On the other side, the neuromorphic behaviours of nanostructured gold (ns-Au)^13,16^ and zirconia (ns-ZrOx) thin films^10,17^ allow the development of functional interfaces for the in materia processing of sensor signals and complex time-series dynamics, on rigid^11,18^ and soft substrates^19,20^ able to respond to environmental and mechanical stimuli.

The combination of nanostructured zirconia and gold (ns-ZrOx/Au) has recently been shown to exhibit memristive behavior with short-term memory, relaxation dynamics, and spike-dependent plasticity.^10,17^ The planar structure of these nanocomposite devices and their biocompatibility make them suitable for the coupling with biological neural networks. Recently, a methodology based on these ns-Au/ZrOx cluster-assembled films and simple statistical analysis methods, has proved the possibility of processing the neural traces recorded by in vivo experiments, without the need for artificial neural networks and preprocessing of the signals.^11^ These nanostructured materials are therefore proposed as a potential solution for the implementation of on-edge devices suitable for functional brain–machine interfaces, featuring low energy and computational consumption, and compatible with in vivo applications.^21,22^

In this study, we investigate the biocompatibility and functional neuroglial interface properties of nanostructured zirconia-based films, focusing on their application as substrates for co-cultures of glial cells and neurons derived from both the central and peripheral nervous systems. Specifically, we compare the cellular responses elicited by nanostructured zirconia versus flat zirconia substrates in primary astrocytes and DRG neuron-glia co-cultures. Our results reveal robust cellular adhesion, survival, and morphological integration of DRG glia and neurons, but also the emergence of distinct calcium signaling profiles that vary by cell type and substrate topography.

These findings underscore the capacity of nanostructured zirconia to modulate calcium signalling mediating neuron-glia communication, with implications for its use in next-generation neural interfaces. The observed substrate-dependent differences in calcium dynamics open the path for the use of material surface properties to selectively influence calcium dependent functional crosstalk between neurons and glia, a hallmark of glia-neuromorphic materials interfaces^2,5,6^.

## MATERIALS AND METHODS

### Nanostructured thin film deposition and characterization

Nanostructured zirconia thin films have been deposited by means of a Supersonic Cluster Beam Deposition (SCBD) apparatus^23^, equipped with a Pulsed Micro Plasma Cluster Source (PMCS) for the production of clusters in the gas phase.^24^ The SCBD set-up schematically consists of two differential pumped vacuum chambers. The first chamber is coupled with a PMCS where the production of zirconia clusters in the gaseous phase is realized through the ablation of a metal target by a discharge plasma after the injection of an inert gas (Ar). The cluster-inert gas mixture is then extracted to form a supersonic seeded beam that impinges on a glass substrate fixed on a sample holder perpendicular to the beam trajectory in the second chamber. The roughness (Rq) of the nanostructured film evolves with thickness, according to simple scaling laws.^25,26^ By recording the film thickness through a quartz microbalance inserted in the deposition chamber, it is possible to finely tune the morphological properties of the film. Flat zirconia samples (Rq⁓0.2 nm), used as control for the evaluation of the role of morphology, have deposited by ion guns on glass substrates.

Topographical maps of the nanostructured zirconia films have been acquired in air using a Multimode Atomic Force Microscopyy (AFM) equipped with a Nanoscope IV controller (BRUKER). Rigid silicon tapping mode cantilevers mounting single crystal silicon tip with nominal radius 5–10 nm and resonance frequency in the range 250–350 kHz have been used. Several 2 μm × 1 μm images were acquired on each sample with scan rate of 1 Hz and 2048 × 512 points. The images were flattened by line-by-line subtraction of first and second order polynomials, in order to remove artifacts due to sample tilt and scanner bow, and subsequently analyzed in Matlab environment.

### Rat cortical astrocytes culture preparation, maintenance, and plating

Primary cultures of astrocytes were prepared from newborn Sprague Dawley rat pups (postnatal days P0–P2) following established protocols. Briefly, neonatal cerebral cortices were dissected to remove the meninges, gently triturated, filtered through a 70 μm cell strainer, and plated in T25 flasks. These flasks contained Dulbecco’s Modified Eagle Medium (DMEM) with GlutaMAX, high glucose, 15% fetal bovine serum (FBS), and penicillin-streptomycin at concentrations of 100 U/mL and 100 μg/mL, respectively. The cultures were maintained in a humidified incubator at 37 °C with 5% CO₂. The medium was refreshed every two days, and flasks were gently shaken as needed to remove microglial cells. After two weeks, when astrocytes reached confluence, cells were detached using 0.25% trypsin-EDTA and re-plated onto Flat-ZrOx, ns-ZrOx, and control substrates. Cells were seeded at a density of 5–7×10³ cells per dish and cultured in medium containing 10% FBS.

### Dorsal root ganglion cell culture preparation

Primary DRG cultures were prepared from postnatal (P7–P21) rats following established protocols^27^. Equal volumes of the DRG cell suspension were applied directly onto the substrates and incubated at 37 °C in 5% CO₂. DRG cells were maintained in DMEM supplemented with 10% FBS and 50 ng/mL nerve growth factor (NGF). Cytosine β-D-arabinofuranoside (AraC, 1.5 mg/mL) was omitted to preserve peripheral glial cell expression.

Cultures were characterized at 3, 5, and 7 DIV using optical and confocal microscopy.

All experiments adhered to Italian laws governing the protection of laboratory animals and were approved by the Bioethical Committee of the University of Bologna and the Ministry of Health (ID 1268, code number 2DBFE.N.HTT, protocol number 360/2017-PR). Oversight was provided by the university’s veterinary commission to ensure animal welfare. Every effort was made to minimize animal usage and reduce suffering throughout the study.

### Fluorescent Diacetate Assay (FDA)

The biocompatibility of astrocyte and DRG neuron-glial cell co-cultures on flat-ZrOx, ns-ZrOx and PDL (used as a control) was assessed at 3 and 7 days in vitro (DIV) for astrocytes and 5 DIV for DRG using the fluorescein diacetate (FDA) assay. Quantitative analysis was conducted by counting all live, FDA-positive (green) cells/total number of nucleai with ImageJ software.

### Differentiation analysis

Astrocyte differentiation induced by interaction with ZrOx substrates was evaluated using morphological criteria. Cells were classified as differentiated when exhibiting an elongated or star-shaped morphology, while polygonal cells were considered undifferentiated. Quantitatively, a cell was defined as differentiated when the length of its major axis was at least twice that of its minor axis.

### Calcium imaging and pharmacology

For experiments involving primary cultures, variations in [Ca2+]i were assessed using calcium microfluorometry with the fluorescent Ca2+ indicator Fluo-4 AM (Life Technologies). Prior to measurements, high-density astrocytes cultured on ZrOx substrates and on PDL were incubated with 2 µM Fluo-4 AM, which was dissolved in a standard bath solution, for 45 minutes at room temperature.

Measurements of [Ca2+]i were conducted using a fluorescence microscope (Nikon Eclipse Ti-S) equipped with a long-distance dry objective (×40) and appropriate filters. The excitation wavelength was set at 450 nm, with a light pulse duration of 200 ms and a sampling rate of 2 Hz. Data acquisition was managed through MetaFluor software (Molecular Devices). Blockers were diluted in standard bath saline to achieve their respective final concentrations and were introduced following rinsing. In the context of in vitro calcium imaging experiments, cells were deemed responsive to stimulation when the maximum change in fluorescence after the stimulus exceeded 0.05 ΔF/F. To analyze the temporal characteristics of [Ca2+]i dynamics, we calculated the average number of peaks by detecting the fluorescence oscillations recorded over time. The onset was calculated at the time point where we could measure the minimal variation (0.02 ΔF/F) in ΔF/F.

### Solution and chemicals

Salts and other high-purity chemicals were sourced from Sigma. For the calcium microfluorometry experiments, the standard bath solution consisted of (mM) 140 NaCl, 4 KCl, 2 MgCl2, 2 CaCl2, 10 HEPES, and 5 glucose, adjusted to a pH of 7.4 with NaOH and an osmolarity of approximately 318 mOsm using mannitol. The calcium-free extracellular saline (NO EXT-Ca2+) was formulated with (mM) 140 NaCl, 4 KCl, 4 MgCl2, 10 HEPES, and 0.5 EGTA, also at a pH of 7.4 with NaOH and adjusted to ∼318 mOsm with mannitol. Stock solution of 2-APB (100 mM) were prepared by dissolving them in methanol and stored at −20 °C.

### Statistical analysis

For calcium imaging in cell cultures, fluorescence time series were manually extracted using both MetaFluor (Molecular Devices) and a dynamic-data-exchange Excel file (Microsoft Office 365). Representative traces and statistical analyses of the extracted data from in vitro calcium imaging experiments were conducted with Microcal Origin 8.5. Bar-dot plots were created using Prism GraphPad 8.0.2. Data comparisons were performed using one-way ANOVA followed by Fisher’s post-test, also utilizing Prism GraphPad 8.0.2. A statistically significant difference was defined as P ≤ 0.05. All data are presented as mean ± s.e.m., with the sample size (n) and number of experiments (N) for each statistical analysis indicated in the figure captions corresponding to specific results. The analysis was based on data collected from at least three independent experiments.

## RESULTS AND DISCUSSION

### Substrate topography and cell density influence astrocyte viability and differentiation

Astrocytes were cultured on three different substrates to evaluate their viability, morphology, and differentiation: Poly-D-lysine (PDL, used as the control) on glass, flat zirconium oxide (flat-ZrOx), and nanostructured zirconium oxide (ns-ZrOx), the latter with a surface roughness of 15 nm. Representative AFM images of the different substrates are reported in figure 1a. Two seeding densities were employed to assess the effect of cell density on astrocyte behaviour: high density (30,000 cells) and low density (15,000 cells). Cell viability and morphology were analyzed using FDA staining, providing key insights into how substrate type and cell density influence astrocyte responses over time.

**Figure 1:**
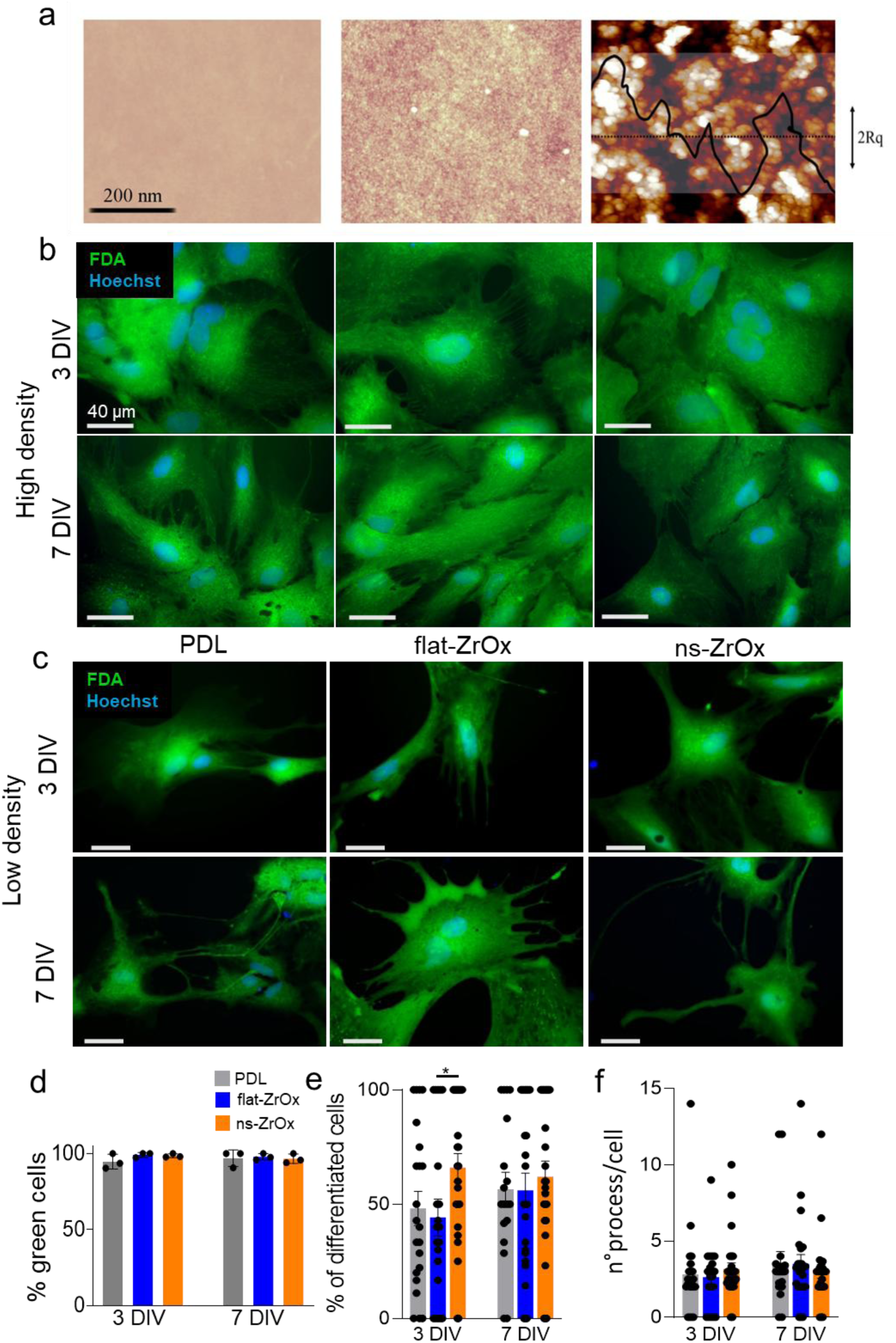
ZrOx substrates support astrocyte viability and morphological differentiation. a) AFM top-view images of PDL, flat-ZrOx and nanostructured (ns) ZrOx substrates; Z-bar scale bar ranges from −20 to 20 nm. b) FDA staining on astrocytes plated on PDL, flat-ZrOx and ns-ZrOx at 3 and 7 DIV at high density. Note, astrocytes are not differentiated in any of the substrates at any of the time points (scale bar: 40µm). c) FDA staining on astrocytes plated on PDL, flat-ZrOx and ns-ZrOx at 3 and 7 DIV at low density (scale bar: 40µm) d) Histogram representing the percentage of green cells (alive cells) plated on PDL at 3 (94,63 ± 2,84) and at 7 DIV (96,88 ± 3,125), on flat-ZrOx at 3 (99,07 ± 0,92) and at 7 DIV (97,78 ± 1,21), and on ns-ZrOx at 3 (98,55 ± 0,72) and at 7 DIV (96,56 ± 1,79). e) Histogram representing the % of differentiated cells in astrocytes plated on PDL at 3 (48,13 ± 7,44) and at 7 DIV (56,54 ± 7,43), on flat-ZrOx at 3 (44,18 ± 8,09) and at 7 DIV (56,12 ± 7,42), and on ns-ZrOx at 3 (65,97 ± 6,19) and at 7 DIV (62,07 ± 6,72). Note, the percentage of differentiated astrocytes on ns-ZrOx is significantly higher with respect to flat-ZrOx (*p= 0,0345). f) Histogram showing the number of processes per cell in astrocytes plated on PDL at 3 (2,83 ± 0,61) and at 7 DIV (3,53 ± 0,77), on flat-ZrOx at 3 (2,62 ± 0,45) and at 7 DIV (3,50 ± 0,59), and on ns-ZrOx at 3 (3,16 ± 0,41) and at 7 DIV (3 ± 0,46).

FDA staining showed that astrocytes plated at both density demonstrated consistent viability across all substrates, as evidenced by the presence of viable cells (green cells) at both 3 days in vitro (DIV) and 7 DIV (fig. 1b,c).

From a morphological point of view the cells plated at high density look polygonal and do not show the presence of processes (fig. 1b). Whereas astrocytes cultured at low density exhibited a pronounced increase in differentiation, as demonstrated by the formation of processes extending from the soma on all substrates (fig. 1c).

Quantitative analysis revealed no significant differences in cell adhesion or viability among the three substrates, as the percentage of viable (green-stained) astrocytes was comparable between the control substrate [PDL (3 DIV: 94,63 ± 2,84. 7 DIV: 96,88 ± 3,125)] and the ZrOx substrates [flat-ZrOx (3 DIV: 99,07 ± 0,92. 7 DIV: 97,78 ± 1,21), ns-ZrOx (3 DIV: 98,55 ± 0,72. 7 DIV: 96,56 ± 1,79)] (fig. 1d). At low cell density, the proportion of differentiated astrocytes showed a slight, though not statistically significant, increase on ns-ZrOx compared to the other substrates [PDL (3 DIV: 48,13 ± 7,44. 7 DIV: 56,54 ± 7,43), flat-ZrOx (3 DIV: 44,18 ± 8,09. 7 DIV: 56,12 ± 7,42), ns-ZrOx (3 DIV: 65,97 ± 6,19. 7 DIV: 62,07 ± 6,72)], with this trend becoming more pronounced at 3 DIV (fig. 1e). In contrast, no significant differences were observed among substrates in the number of cellular processes per cell [PDL (3 DIV: 2,83 ± 0,61. 7 DIV: 3,53 ± 0,77) flat-ZrOx (3 DIV: 2,62 ± 0,45. 7 DIV: 3,50 ± 0,59), ns-ZrOx 3 DIV: 3,16 ± 0,41. 7 DIV: 3 ± 0,46)] (fig. 1f).

Collectively, these data indicate that from a morphological standpoint, either astrocytes plated on PDL and on ZrOx substrates at high density maintained an undifferentiated state, with an almost complete absence of differentiated cells at both time points (data not shown). This suggests that high-density conditions may inhibit the initiation of differentiation, potentially due to limited space for cell extension or altered intercellular signalling in crowded environments ^28^.

The reduced cell-cell contact, and increased availability of substrate surface may promote signalling pathways and cytoskeletal remodelling necessary for process extension and maturation. The results suggest that substrate chemistry by itself does not significantly alter astrocyte viability but may interact with cell density to influence differentiation. Specifically, while flat-ZrOx and ns-ZrOx supported similar viability to the control, the low-density seeded cell condition appeared more permissive for differentiation across all substrates. These findings are in agreement with previous reports indicating that cell seeding density modulate cell morphology and functional state^29,30^, providing valuable insights for biomaterial development and neural tissue engineering applications. Substrate topography is well known to influence key cellular processes, including migration, adhesion, and polarization; in stem cells, it has also been shown to play a crucial role in guiding differentiation^31,32^. Similarly, studies on primary astrocytes have demonstrated that nanostructured surfaces can enhance their differentiation rate^33,34^. Astrocytes differentiation on nanostructured surfaces is frequently accompanied by an increase in calcium signalling^35^. Activation of calcium dependent signaling pathways, such as RHO, TGF-β, or FAK can drive cells toward distinct differentiation states^36–38^; notably, these pathways are tightly interconnected with canonical mechanotransduction mechanisms, which are themselves responsive to substrate nanotopography^31^.

### Substrate nanotopography differentially modulates intracellular calcium signaling pathways in astrocytes

Given the importance of calcium signalling on astrocytes and the functional response of astrocytes to different substrates, we performed calcium imaging on cells cultured on PDL, flat-ZrOx and ns-ZrOx. This analysis revealed significant differences in intracellular calcium ([Ca²⁺]i) dynamics between the control and ZrOx substrates, highlighting the influence of substrate material and structure on astrocyte behaviour.

Figure 2b shows the temporal changes in calcium intensity, revealing substrate-driven calcium dynamics. Astrocytes cultured on ns-ZrOx exhibited a markedly stronger maximal calcium amplitude (ΔF/F) (0,30 ± 0,036, n = 55, N = 10) compared to those on the control substrate (0,19 ± 0,016, n = 78, N = 10) (fig. 2c), accompanied by a significantly shorter time onset (ns-ZrOx: 111,1 ± 9,49, n = 55, N = 10. PDL: 166,5 ± 8,47, n = 78, N = 10) (fig. 2d) and time to peak (ns-ZrOx: 213 ± 10,84, n = 55, N = 10. PDL: 247,6 ± 5,53, n = 78, N = 10) (fig. 2e).

**Figure 2:**
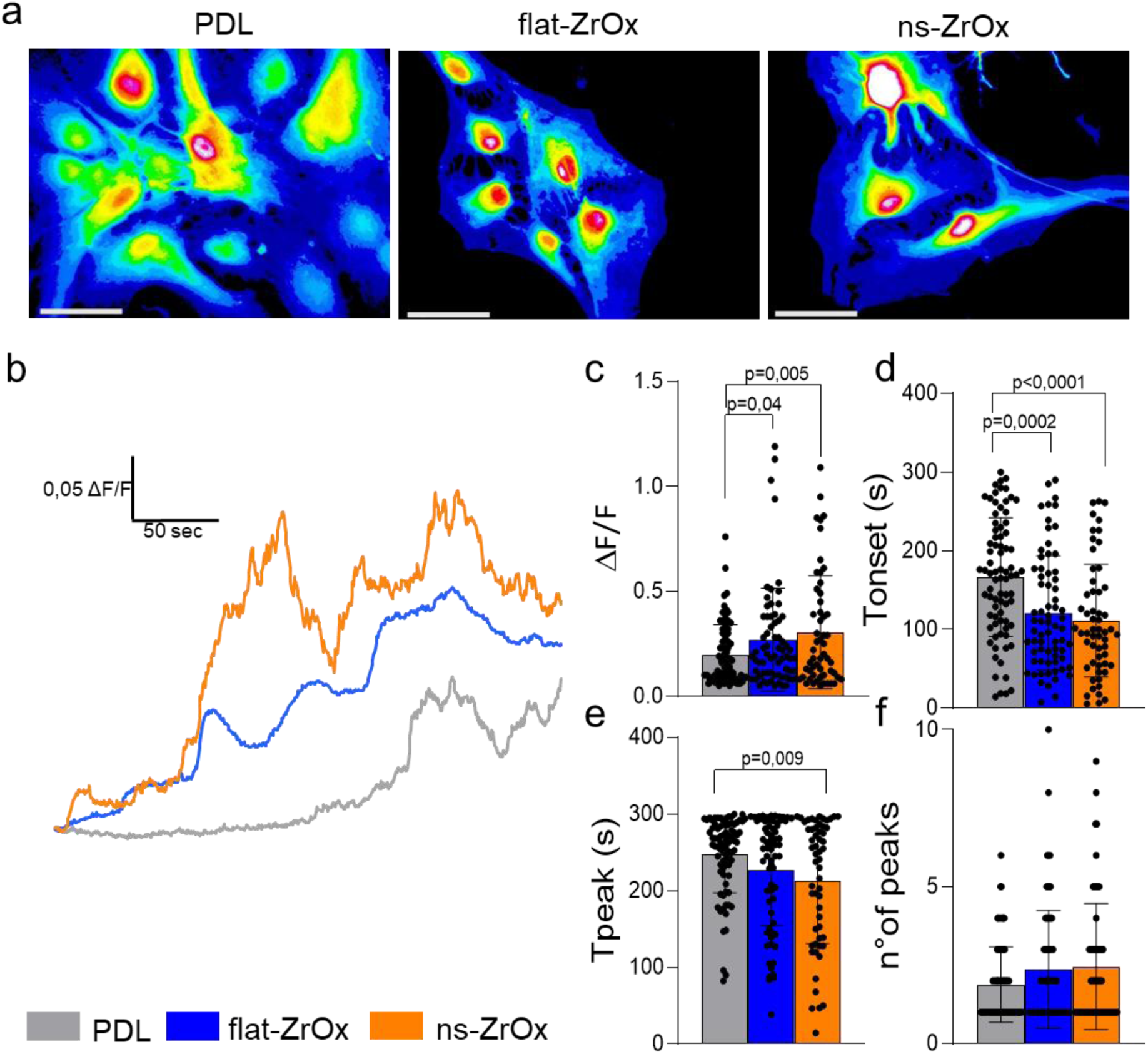
ZrOx modulates spontaneous calcium dynamics in astrocytes. a) Representative images illustrating the spatial distribution of calcium waves in astrocytes grown on PDL, flat-ZrOx, and ns-ZrOx surfaces (scale bar: 40µm). b) Representative traces of fluorescence intensity variation (ΔF/F) over time of the astrocytes plated on the different substrates. c) Histogram showing maximum fluorescence intensity values in responding astrocytes cultured on PDL (0,19 ± 0,016, n = 78, N = 10), flat-ZrOx (0,26 ± 0,029, n = 68, N = 10), and ns-ZrOx (0,30 ± 0,036, n = 55, N = 10). Astrocytes on PDL showed significantly lower maximum fluorescence intensity compared to flat-ZrOx (*p = 0.04) and ns-ZrOx (**p = 0.005). d) Histogram showing the time onset in responding astrocytes cultured on PDL (166,5 ± 8,47, n = 78, N = 10), flat-ZrOx (120,7 ± 8,75, n = 68, N = 10), and ns-ZrOx (111,1 ± 9,49, n = 55, N = 10). Astrocytes on PDL showed significantly slower response compared to flat-ZrOx (***p = 0.0002) and ns-ZrOx (****p < 0.0001). e) Histogram showing the time to peak in responding astrocytes cultured on PDL (247,6 ± 5,53, n = 78, N = 10), flat-ZrOx (226,8 ± 8,71, n = 68, N = 10), and ns-ZrOx (213 ± 10,84, n = 55, N = 10). Astrocytes on PDL showed significantly slower time to peak compared to ns-ZrOx (**p =0.009). f) Histogram showing the number of peaks in responding astrocytes cultured on PDL (1,88 ± 0,13, n = 78, N = 10), flat-ZrOx (2,36 ± 0,22, n = 68, N = 10), and ns-ZrOx (2,45 ± 0,26, n = 55, N = 10).

These findings indicate that the nanostructured surface not only amplified the calcium response but also accelerated its kinetics, suggesting a more rapid and intense cellular activation. Similarly, astrocytes on flat-ZrOx displayed a significantly shorter time onset (120,7 ± 8,75, n = 68, N = 10) compared to the control substrate, although the differences on maximal amplitude (0,26 ± 0,029, n = 68, N = 10) and time to peak (226,8 ± 8,71, n = 68, N = 10) were less pronounced than on ns-ZrOx. Interestingly, the number of calcium peaks, indicative of the frequency of calcium oscillations, did not differ significantly across the three substrates (PDL: 1,88 ± 0,13, n = 78, N = 10. Flat-ZrOx: 2,36 ± 0,22, n = 68, N = 10. Ns-ZrOx: 2,45 ± 0,26, n = 55, N = 10) (fig. 2f). This suggests that while the material and nanostructured surface modulate the intensity and speed of calcium signaling, they do not alter the overall pattern of calcium dynamics. These results demonstrate that substrate chemistry do not affect cell viability, substrate chemical composition and its topography directly influence the functionality of astrocytes, namely their calcium signalling. Specifically, the nanostructured ns-ZrOx surface enhances both the amplitude and shorten the onset of the calcium response, suggesting that nanoscale features may optimize substrate-cell interactions, potentially by affecting ion channel activity, receptor clustering, or mechanosensitive pathways. These findings provide valuable insights into how material morphology can be leveraged to modulate astrocyte function, with implications for biomaterial development and neural tissue engineering.

We next applied selective pharmacological approaches to investigate the contribution of calcium influx and internal stores to the substrate-dependent differences in astrocytes’ calcium signalling.

2-Aminoethyl Diphenylborinate (2APB), a cell-permeable inhibitor of inositol 1,4,5-trisphosphate (Ins(1,4,5)P3 or IP3)-induced Ca²⁺ release, was used to assess the role of IP3 signaling in astrocyte calcium dynamics across different substrates.

The calcium-free extracellular saline (0Ca²⁺) was employed to eliminate extracellular calcium flux, thereby enabling the exclusive observation of intracellular calcium dynamics. This approach isolates the contributions of internal calcium stores and pathways, providing a clearer understanding of intracellular calcium signaling mechanisms.

Figure 3a shows representative calcium traces in astrocytes cultured on distinct substrates, following both IP₃-dependent signaling inhibition and extracellular calcium removal.

**Figure 3:**
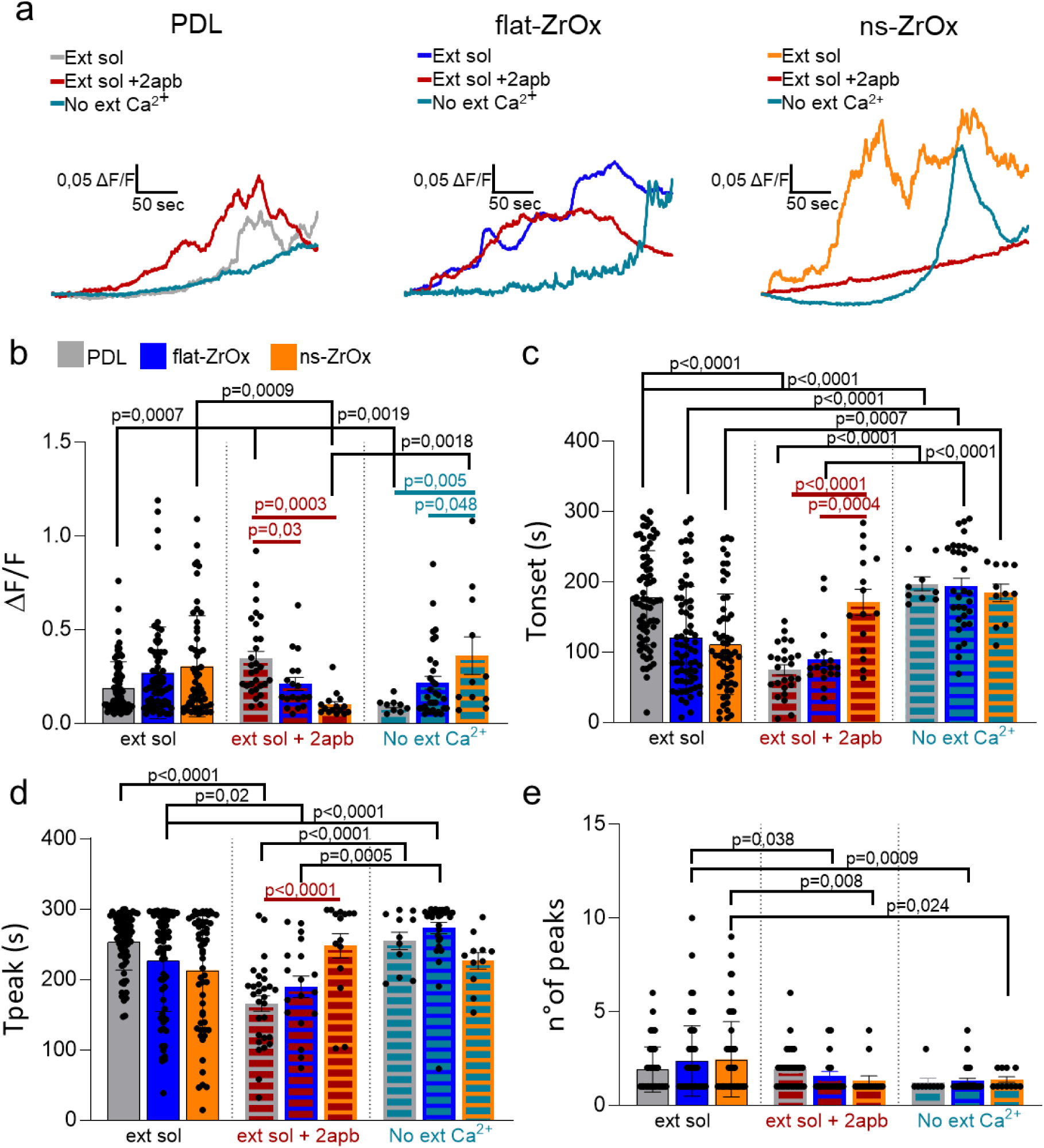
Astrocytes calcium signaling on ZrOx depends on both intracellular release and extracellular Ca^2+^ entry. a) Representative traces showing ΔF/F fluorescence intensity changes over time in astrocytes on all substrates, comparing control conditions with IP₃ pathway inhibition and removal of external Ca²⁺. b) Histogram showing maximum fluorescence intensity values in responding astrocytes cultured on PDL, flat-ZrOx, and ns-ZrOx in control conditions, following IP₃ pathway inhibition [PDL (0,34 ± 0,038, n = 29, N = 3), flat-ZrOx (0,20 ± 0,034, n = 18, N = 3), and Ns-ZrOx (0,099 ± 0,016, n = 15, N = 3)] and removal of external Ca²⁺ [PDL (0,09 ± 0,011, n = 9, N = 3), Flat-ZrOx (0,21 ± 0,034, n = 32, N = 3), and ns-ZrOx (0,36 ± 0,1, n = 11, N = 3)]. c) Histogram showing the time onset in responding astrocytes cultured on PDL, flat-ZrOx, and ns-ZrOx in control conditions, following IP₃ pathway inhibition [PDL (74,62 ± 7,25, n = 29, N = 3), flat-ZrOx (90,06 ± 10,30, n = 18, N = 3), and ns-ZrOx (170,9 ± 18,43, n = 15, N = 3)] and removal of external Ca²⁺ [PDL (197,2 ± 9,83, n = 9, N = 3), flat-ZrOx (194,3 ± 10,83, n = 32, N = 3), and ns-ZrOx (184,1 ± 12,44, n = 11, N = 3)]. d) Histogram showing the time to peak in responding astrocytes cultured on PDL, flat-ZrOx, and ns-ZrOx in control conditions, following IP₃ pathway inhibition [PDL (165,8 ± 10,94, n = 29, N = 3), flat-ZrOx (189,5 ± 15,11, n = 18, N = 3), and ns-ZrOx (247,9 ± 17,47, n = 15, N = 3)] and removal of external Ca²⁺ [PDL (254,7 ± 12,49, n = 9, N = 3), flat-ZrOx (273,3 ± 7,99, n = 32, N = 3), and ns-ZrOx (226,3 ± 11,75, n = 11, N = 3)]. e) Histogram showing the number of peaks in responding astrocytes cultured on PDL, flat-ZrOx, and ns-ZrOx in control conditions, following IP₃ pathway inhibition [PDL (1,93 ± 0,21, n = 29, N = 3), flat-ZrOx (1,55 ± 0,24, n = 18, N = 3), and fs-ZrOx (1,33 ± 0,23, n = 15, N = 3)] and removal of external Ca²⁺ [PDL (1,22 ± 0,22, n = 9, N = 3), flat-ZrOx (1,31 ± 0,12, n = 32, N = 3), and ns-ZrOx (1,36 ± 0,15, n = 11, N = 3)]. p values are displayed on the figure.

The data, illustrated in Figure 3, reveal significant alterations in astrocyte calcium activity upon 2APB treatment, particularly for cells cultured on ns-ZrOx. Astrocytes on ns-ZrOx exhibited a pronounced reduction in maximal calcium amplitude (ΔF/F) (ns-ZrOx: 0,099 ± 0,016, n = 15, N = 3) (fig. 3b), indicating diminished release from intracellular calcium stores. Additionally, the time onset of calcium transients was significantly increased (ns-ZrOx: 170,9 ± 18,43, n = 15, N = 3), demonstrating a delay in the initiation of calcium signaling. The frequency of calcium peaks, which reflects the oscillatory behavior of calcium dynamics, was also markedly reduced in astrocytes on ns-ZrOx (Ns-ZrOx: 1,33 ± 0,23, n = 15, N = 3). These findings suggest that the nanostructured surface amplifies IP3-dependent calcium signaling under normal conditions, making it particularly susceptible to disruption by 2APB. The observed changes were less evident on other substrates, underscoring the unique influence of the ns-ZrOx surface on IP3-mediated calcium dynamics.

Interestingly, the removal of extracellular Ca²⁺ did not affect the maximal calcium amplitude (ΔF/F) in ZrOx (Flat-ZrOx: 0,21 ± 0,034, n = 32, N = 3. Ns-ZrOx 0,36 ± 0,1, n = 11, N = 3) (fig. 3b), as the values remained comparable to those observed under normal conditions. However, the absence of extracellular calcium significantly influenced other critical aspects of calcium signaling. The time to onset of the calcium response was markedly prolonged (PDL: 197,2 ± 9,83, n = 9, N = 3. Flat-ZrOx: 194,3 ± 10,83, n = 32, N = 3. Ns-ZrOx (184,1 ± 12,44, n = 11, N = 3) (fig. 3c), indicating that extracellular calcium may play a role in accelerating the initiation of intracellular calcium release. Additionally, the number of calcium peaks was significantly reduced on ZrOx substrates (Flat-ZrOx: 1,31 ± 0,12, n = 32, N = 3. Ns-ZrOx: 1,36 ± 0,15, n = 11, N = 3) (fig. 3e), suggesting that extracellular calcium contributes to the frequency modulation of calcium transients. These findings imply that while the magnitude of calcium signaling can be sustained through intracellular stores, extracellular calcium is essential for fine-tuning the temporal and dynamic features of calcium signaling pathways.

The ability of ns-ZrOx to amplify intracellular calcium responses suggests that the nanostructured interface enhances the mechanosensitivity of astrocyte membranes, possibly by reorganizing integrin clusters, focal adhesions, or associated ion channels such as TRP or Piezo family members^39,40^.

The application of 2APB, an IP₃ receptor antagonist, revealed that the enhanced calcium signaling on ns-ZrOx is strongly dependent on IP₃-mediated intracellular calcium release. The pronounced sensitivity of astrocytes to IP₃R inhibition on the nanostructured surface indicates that the substrate actively promotes ER-plasma membrane communication or potentiates phosphoinositide signaling pathways^1,41^. Interestingly, in calcium-free conditions, astrocytes maintained similar signal amplitudes but displayed delayed response kinetics and reduced oscillation frequency, supporting the hypothesis that extracellular calcium influx contributes primarily to the temporal modulation rather than the magnitude of intracellular calcium dynamics.

### Zirconia-based substrates demonstrate high biocompatibility with DRG neuron–glia co-cultures and drive their functional behaiviour

To extend our investigation of zirconia-based substrates beyond the CNS and into the PNS, we established a co-culture model of DRG neurons and glial cells. DRG cultures represent a well-established and physiologically relevant system for studying peripheral sensory neurons and their interactions with associated glia, and they serve as a powerful in vitro model to explore sensory system development, injury responses, and neuroregenerative mechanisms^42^.

As illustrated in Figure 4a,b, all tested ZrOx substrates exhibited excellent biocompatibility with DRG co-cultures. FDA images show the presence of green (alive) cells with different morphological features (fig. 4a). The quantitative analysis of cell survival indicates no significant impairment of cell adhesion across the groups (PDL: 95.70 ± 2.22. Flat-ZrOx: 99.37 ± 0.46. Ns-ZrOx: 98.12 ± 1.22) (fig. 4b). Importantly, DRG neurons can be clearly identified by their characteristic spherical or ovoid somata, often grouped in clusters, which is consistent with their native morphology in vivo.

**Figure 4:**
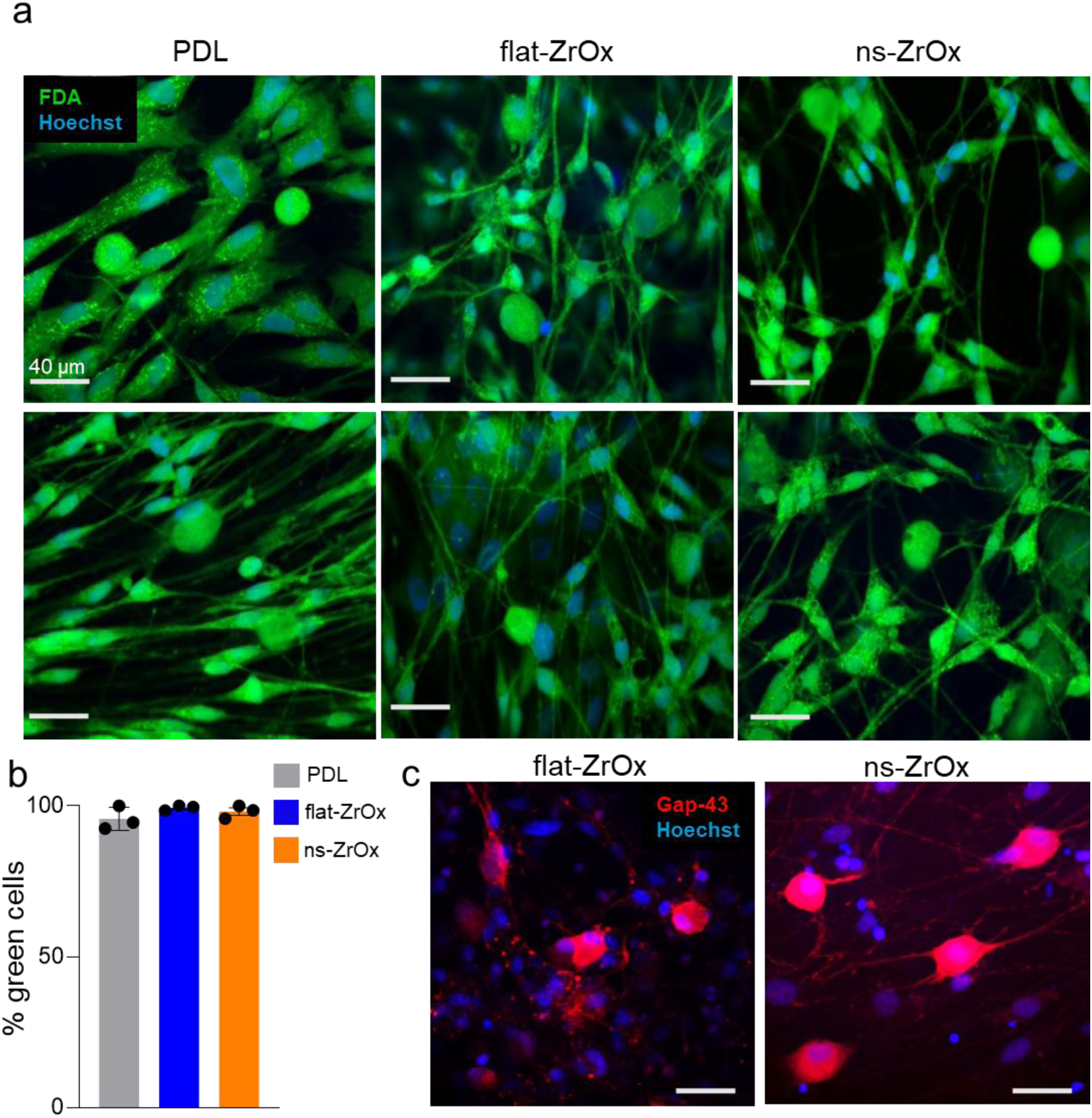
ZrOx support DRG neuron-glia viability. a) FDA staining of DRG neuron–glia co-cultures plated on PDL, flat-ZrOx, and ns-ZrOx substrates at 5 DIV (scale bar: 40µm). b) Histogram showing the percentage of viable (green) cells on PDL (95.70 ± 2.22), Flat-ZrOx (99.37 ± 0.46), and ns-ZrOx (98.12 ± 1.22). c) Immunofluorescence images of Gap-43–positive cells. Rounded Gap-43–positive cells are likely neuron (scale bar: 40µm).

This neuronal identity was further validated through immunocytochemical staining for GAP-43 (Growth Associated Protein-43), a well-established marker of axonal growth and regeneration that is particularly prominent in developing or regenerating DRG neurons (fig. 4c)^43^.

The glial cells, which surround and support the neurons in culture, also showed typical elongated morphologies and close association with neuronal somata and neurites, consistent with functional neuron-glia interactions. Together, these results confirm that zirconia substrates support healthy co-culture conditions for DRG-derived cells, validating their applicability in PNS-targeted neuroengineering approaches and supporting their potential use in biointerfaces for peripheral nerve repair or stimulation.

To investigate how the substrate morphology and chemistry influence cellular excitability and signaling, we performed a detailed analysis of calcium dynamics in DRG co-cultures seeded on the three different substrates. Calcium transients were recorded in both neuronal and glial populations under the previous distinct conditions: (i) control physiological solution, (ii) following the administration 2APB, and (iii) in calcium-free extracellular medium. Figure 5 presents calcium responses of DRG neurons and glial cells on three different substrates under control conditions. Representative traces in Figure 5b reveal the substrate-specific temporal dynamics of calcium signaling in both cell types. Notably, glial cells on both flat-ZrOx (0,32 ± 0,026, n = 98, N=10) and ns-ZrOx (0,34 ± 0,028, n = 97, N=10) showed significantly higher ΔF/F amplitudes compared to those cultured on PDL (0,23 ± 0,023, n = 83, N=10) (fig. 5c), suggesting that zirconia substrates enhance glial responsiveness. Among the zirconia variants, ns-ZrOx induced also in co-culture the shortest onset time (Flat-ZrOx: 103,2 ± 6,15, n = 68, N = 10. Ns-ZrOx: 85,46 ± 5,83, n = 97, N=10) (fig. 5d), indicating faster calcium entry and/or mobilization from intracellular stores. This accelerated response may reflect enhanced mechanosensitivity or improved interface-mediated signaling due to the nanoscale topography. While we do not observe any difference in the time to peak and the number of peak in glial calcium signalling among the substrates (fig. 5e,f)

**Figure 5:**
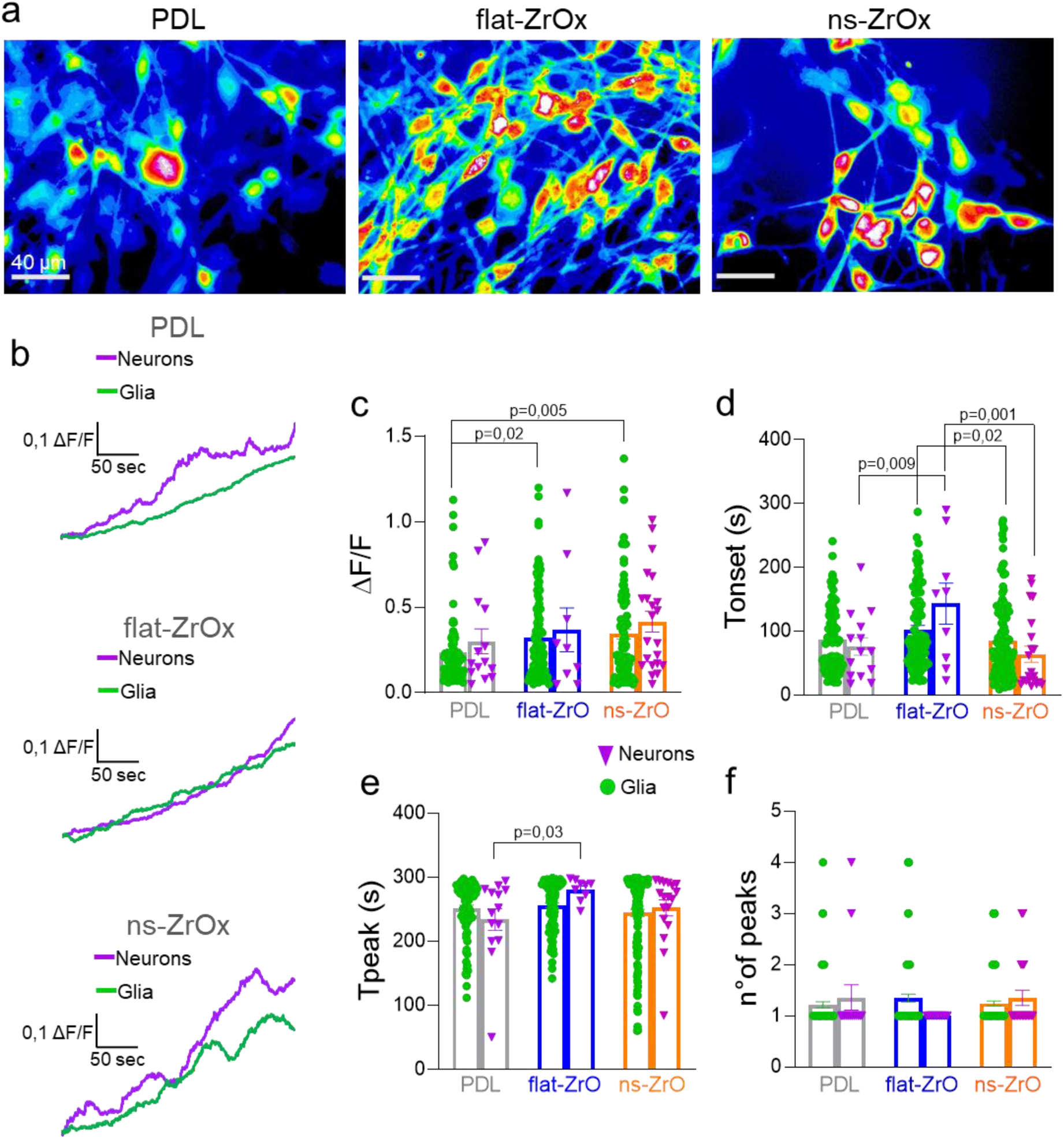
ZrOx enhances calcium signaling in DRG neuron-glia co-cultures. a) Representative images illustrating the spatial distribution of calcium waves in DRG neuron-glia co-culture grown on PDL, Flat-ZrOx, and Ns-ZrOx surfaces (scale bar: 40µm). b) Representative traces of fluorescence intensity variation (ΔF/F) over time of DRG neuron and glia plated on all the substrates. c) Histogram showing maximum fluorescence intensity values in responding astrocytes cultured on PDL (glia: 0,23 ± 0,023, n = 83, neuron: 0,30 ± 0,07, n = 14 N = 10), Flat-ZrOx (glia: 0,32 ± 0,026, n = 68, neuron: 0,37 ± 0,12, n = 9, N = 10), and Ns-ZrOx (glia: 0,34 ± 0,028, n = 97, neuron: 0,41 ± 0,06, n = 22, N = 10). d) Histogram showing the time onset in responding DRG neuron and glia cultured on PDL (glia: 87,15 ± 5,63, n = 83, neuron: 76,32 ± 13,91, n = 14 N = 10), Flat-ZrOx (glia: 103,2 ± 6,15, n = 68, N = 10, neuron: 143,4 ± 32,2, n = 9 N = 10), and Ns-ZrOx (glia: 85,46 ± 5,83, n = 97, neuron: 64,21 ± 12,6, n = 21, N = 10). e) Histogram showing the time to peak in responding DRG neuron and glia cultured on PDL (glia: 251,9 ± 4,90, n = 83, neuron: 234,2 ± 17,08, n = 14 N = 10), Flat-ZrOx (glia: 256 ± 3,91, n = 68,N = 10, neuron: 280,8 ± 5,56, n = 9 N = 10), and Ns-ZrOx (glia: 244,8 ± 5,42, n = 97, neuron: 252,5 ± 12,47, n = 21, N = 10). f) Histogram showing the number of peak in responding DRG neuron and glia cultured on PDL (glia: 1,27 ± 0,06, n = 83, neuron: 1,35 ± 0,24, n = 14 N = 10), Flat-ZrOx (glia: 1,34 ± 0,075, n = 68, N = 10, neuron: 1 ± 0, n = 9 N = 10), and Ns-ZrOx (glia: 1,24 ± 0,04, n = 97, neuron: 1,35 ± 0,15, n = 21, N = 10). p values are displayed on the figure.

In neurons, cultures on ns-ZrOx exhibited a trend toward stronger calcium responses, although differences in ΔF/F amplitude across substrates did not reach statistical significance (fig. 5c). Nevertheless, the consistently higher baseline activity (0,41 ± 0,06, n = 22, N = 10) and shorter time of onset (64,21 ± 12,6, n = 21, N = 10) observed on ns-ZrOx relative to flat-ZrOx (ΔF/F amplitude: 0,37 ± 0,12, n = 9, N = 10. Time onset: 143,4 ± 32,2, n = 9 N = 10) support the hypothesis that nanostructuring enhances neuronal excitability or responsiveness (fig 5c,d). Neurons plated on flat-ZrOx displayed a significantly increased time to peak (280,8 ± 5,56, n = 9 N = 10) with respect PDL (234,2 ± 17,08, n = 14 N = 10), while ns-ZrOx did not show any significant difference (252,5 ± 12,47, n = 21, N = 10) (fig. 5e). The number of peak was similar across the groups [PDL (glia: 1,27 ± 0,06, n = 83, neuron: 1,35 ± 0,24, n = 14 N = 10), flat-ZrOx (glia: 1,34 ± 0,075, n = 68, N = 10, neuron: 1 ± 0, n = 9 N = 10), ns-ZrOx (glia: 1,24 ± 0,04, n = 97, neuron: 1,35 ± 0,15, n = 21, N = 10)] (fig. 5f). These results suggest that nanostructured zirconia not only supports robust cell viability but also actively modulates functional calcium signaling, especially in glial populations.

Given the central role of calcium in neuron-glia communication and plasticity these findings underscore the glio-neuromorphic potential of ns-ZrOx substrates. Figure 6 show the percentage of responsive DRG neurons and glia plated on the three substrates in control condition and following the treatment with 2APB and in absence of extracellular calcium. Following treatment with 2APB, we observed distinct substrate-dependent alterations in calcium signaling within both neuronal and glial populations of the DRG co-cultures. Specifically, neuronal calcium activity was completely abolished on PDL (fig. 6a) and flat-ZrOx (fig. 6b), whereas ns-ZrOx (23,06 ± 11%; N = 4) (fig. 6c) preserved residual calcium signaling, indicating a partial resistance to IP₃R inhibition or the possible contribution of alternative calcium sources, such as mechanosensitive pathways supported by the nanostructured interface.

**Figure 6:**
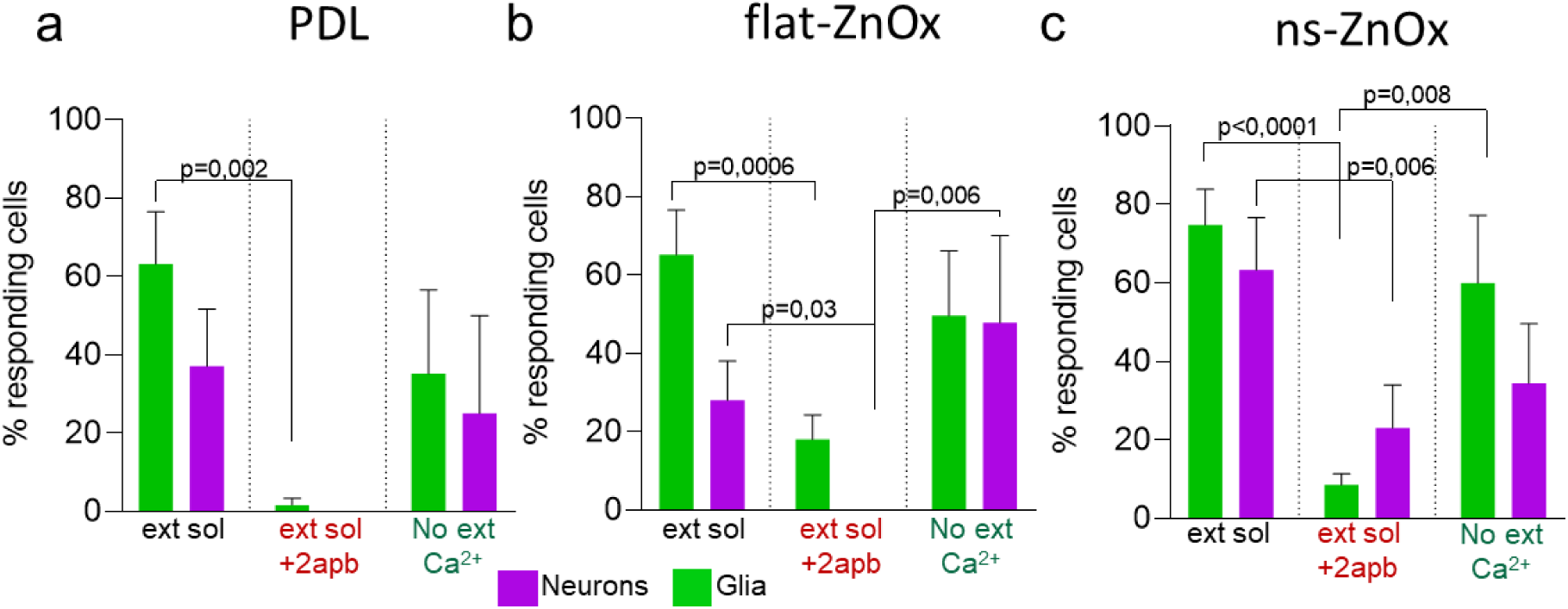
Modulation of DRG neuron and glia responsiveness by internal and external calcium pathways. a) Histogram showing the percentage of responding DRG neurons and glial cells cultured on PDL under different experimental conditions. Data are presented as mean ± SEM for control (glia: 63.23 ± 13.34%; neurons: 37.04 ± 14.7%; N = 10), following 2-APB treatment (glia: 1.67 ± 1.67%; neurons: 0 ± 0%; N = 4), and after extracellular calcium depletion (glia: 35.21 ± 21.35%; neurons: 25 ± 25%; N = 4). b) Histogram showing the percentage of responding DRG neurons and glial cells cultured on flat-ZrOx under different experimental conditions. Data are presented as mean ± SEM for control (glia: 65.23 ± 11.38%; neurons: 28.00 ± 10.6%; N = 10), following 2-APB treatment (glia: 18 ± 6,23%; neurons: 0 ± 0%; N = 4), and after extracellular calcium depletion (glia: 49,66 ± 16,59%; neurons: 47,92 ± 22,15%; N = 4) c) Histogram showing the percentage of responding DRG neurons and glial cells cultured on ns-ZrOx under different experimental conditions. Data are presented as mean ± SEM for control (glia: 74,77 ± 9,14%; neurons: 63,33 ± 13,33%; N = 10), following 2-APB treatment (glia: 8,53 ± 2,86%; neurons: 23,06 ± 11%; N = 4), and after extracellular calcium depletion (glia: 59,95 ± 17,25%; neurons: 34,43 ± 15,20%; N = 4).

Glial cells in the DRG preparation exhibited a significant reduction in the percentage of responding cells across all substrates (PDL: 1.67 ± 1.67%, N=10. flat-ZrOx: 18 ± 6,23%, N=10. ns-ZrOx: 8,53 ± 2,86%) following 2APB treatment, with respect in control conditions (PDL: 63.23 ± 13.34% N=10. flat-ZrOx: 65.23 ± 11.38%, N=10. ns-ZrOx: 74,77 ± 9,14%, N=10), suggesting a strong dependence on IP₃R-mediated signaling. Whereas in absence of calcium in the extracellular solution, a good percentage of both glial and neuronal cells maintained the activity [PDL: glia: 35.21 ± 21.35%; neurons: 25 ± 25%; N = 4. Flat-ZrOx: (glia: 49,66 ± 16,59%; neurons: 47,92 ± 22,15%; N = 4). ZrOx-(glia: 59,95 ± 17,25%; neurons: 34,43 ± 15,20%; N = 4)] (Fig. 6).

As shown in Figure 7, inhibition of IP₃ signaling and depletion of extracellular calcium significantly alter calcium dynamics in glial cells across all three substrates.

**Figure 7:**
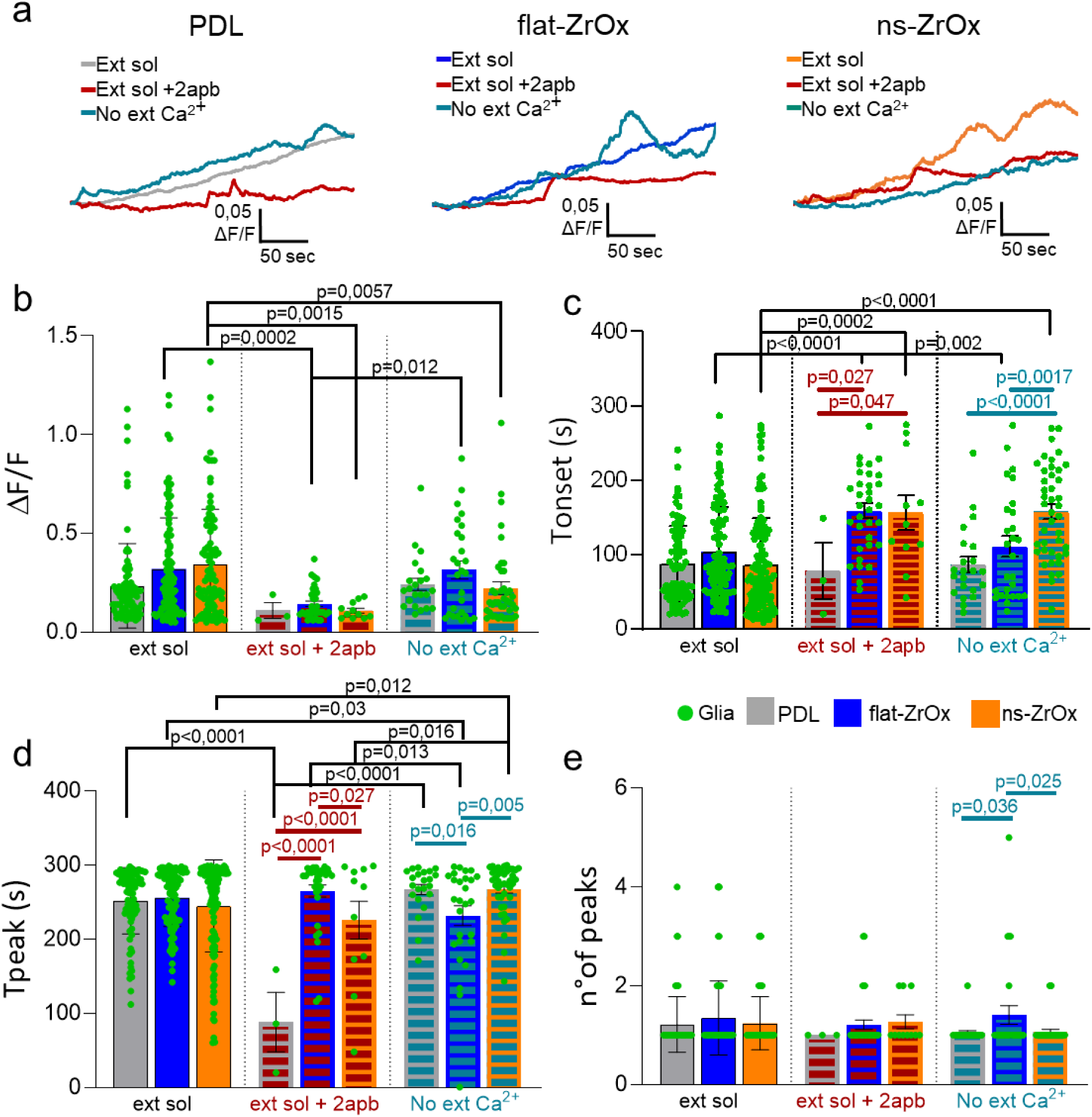
Internal and external calcium contributions to calcium dynamics in DRG glial cells on ZrOx. a) Representative traces of fluorescence intensity variation (ΔF/F) over time of DRG glia plated on all the substrates in control condition and following treatment with 2apb and removal of external Ca²⁺. b) Histogram showing maximum fluorescence intensity values in responding DRG glial cells cultured on PDL, flat-ZrOx, and ns-ZrOx in control conditions, following IP₃ pathway inhibition [PDL (0,11 ± 0,04, n = 3, N = 3), flat-ZrOx (0,14 ± 0,014, n = 33, N = 3), and ns-ZrOx (0,10 ± 0,012, n = 11, N = 3)] and removal of external Ca²⁺ [PDL (0,24 ± 0,03, n = 22, N = 3), Flat-ZrOx (0,24 ± 0,045, n = 27, N = 3), and Ns-ZrOx (0,22 ± 0,03, n = 41, N = 3)]. c) Histogram showing the time onset in responding DRG glial cells cultured on PDL, Flat-ZrOx, and Ns-ZrOx in control conditions, following IP₃ pathway inhibition [PDL (78 ± 37,80, n = 3, N = 3), Flat-ZrOx (158,9 ± 9,53, n = 33, N = 3), and Ns-ZrOx (156,4 ± 23,49, n = 11, N = 3)] and removal of external Ca²⁺ [PDL (86,68 ± 10,7, n = 22, N = 3), Flat-ZrOx (110,4 ± 13,91, n = 27, N = 3), and Ns-ZrOx (157,5 ± 9,58, n = 42, N = 3)]. d) Histogram showing the time to peak in responding DRG glial cells cultured on PDL, Flat-ZrOx, and Ns-ZrOx in control conditions, following IP₃ pathway inhibition [PDL (88,33 ± 40,14, n = 3, N = 3), Flat-ZrOx (265,2 ± 8,19, n = 33, N = 3), and Ns-ZrOx (225,7 ± 25,54, n = 11, N = 3)] and removal of external Ca²⁺ [PDL (267,2 ± 6,88, n = 22, N = 3), Flat-ZrOx (232,1 ± 13,34, n = 27, N = 3), and Ns-ZrOx (267,7 ± 5,31, n = 41, N = 3)]. e) Histogram showing the number of peaks in responding DRG glial cells cultured on PDL, Flat-ZrOx, and Ns-ZrOx in control conditions, following IP₃ pathway inhibition [PDL (1 ± 0, n = 3, N = 3), Flat-ZrOx (1,21 ± 0,09, n = 33, N = 3), and Ns-ZrOx (1,27 ± 0,14, n = 11, N = 3)] and removal of external Ca²⁺ [PDL (1,04 ± 0,04, n = 22, N = 3), Flat-ZrOx (1,40 ± 0,18, n = 27, N = 3), and Ns-ZrOx (1,07 ± 0,04, n = 41, N = 3)]. p values are displayed on the figure.

Following IP₃R blockage we observed that on ns-ZrOx, the ΔF/F amplitude in glial cells was significantly reduced (0,10 ± 0,012, n = 11, N = 3) (fig. 7b), and the time of onset was markedly increased (156,4 ± 23,49, n = 11, N = 3) (fig. 7c), while time to peak (225,7 ± 25,54, n = 11, N = 3) (fig. 7d) and number of peak (1,27 ± 0,14, n = 11, N = 3) (fig. 7e) did not differ from the control media. A similar trend was observed on flat-ZrOx, where glial responses were also diminished in both amplitude (0,14 ± 0,014, n = 33, N = 3) (fig. 7b) and temporal kinetics (158,9 ± 9,53, n = 33, N = 3) (fig. 7c).

Interestingly, glial cells cultured on PDL did not exhibit a statistically significant reduction in calcium response amplitude (0.11 ± 0.04, n = 3, N = 3) or in response timing (78 ± 37.80, n = 3, N = 3) following 2APB treatment, suggesting that calcium dynamics on this conventional substrate may involve additional or compensatory mechanisms, or reflect lower basal IP₃R engagement.

Across all conditions and substrates, both the time of onset and time to peak of calcium transients were increased following 2APB administration, with the most pronounced delays observed on ns-ZrOx (Fig.7). These findings highlight a substrate-specific modulation of intracellular calcium signaling pathways and further reinforce the notion that ns-ZrOx differentially interfaces with glial and neuronal physiology.

In the absence of extracellular calcium, we observed substrate-dependent alterations in intracellular calcium dynamics within both glial cells and neurons cultured on ZrOx substrates. Glial cells plated on ns-ZrOx exhibited a marked reduction in the ΔF/F amplitude (0,22 ± 0,03, n = 41, N = 3) (fig. 7b), accompanied by a significant increase in both the time of onset (157,5 ± 9,58, n = 42, N = 3) (fig. 7c) and the time to peak (267,7 ± 5,31, n = 41, N = 3) of the calcium transient (fig. 7d). These findings indicate a strong dependence on extracellular calcium influx for initiating and shaping calcium signals in glia on ns-ZrOx, possibly due to enhanced membrane interaction or modulation of mechanosensitive or store-operated calcium channels by the nanostructured surface.

Conversely, glial cells on flat-ZrOx maintained a preserved ΔF/F amplitude (0,24 ± 0,045, n = 27, N = 3) (fig. 7b), and notably, the time to peak was shorter (232,1 ± 13,34, n = 27, N = 3) (fig. 7d), suggesting a more stable or sustained reliance on intracellular calcium stores, and potentially a reduced sensitivity to the removal of extracellular calcium.

In figure 8 we show the properties of neuronal calcium dynamics plated on all the substrates. In the neuronal population, the magnitude of the calcium response (ΔF/F) was significantly reduced across both zirconia substrates (flat-ZrOx: 0,18 ± 0,040, n = 7, N = 3. and ns-ZrOx: 0,23 ± 0,08, n = 6, N = 3) (fig. 8b), indicating a critical role of extracellular calcium influx in neuronal calcium signaling regardless of surface topography. However, only neurons on ns-ZrOx exhibited a significantly delayed time of onset (159,8 ± 24,8, n = 6, N = 3) (fig. 8c), suggesting that nanostructuring the surface may affect the kinetics of calcium mobilization, possibly by altering membrane potential, receptor distribution, or the availability of residual intracellular stores. Time to peak results significantly reduced in DRG neurons plated on flat-ZrOx (233,9 ± 12,31, n = 7, N = 3) (fig. 8d), while the number of peaks result stable in all the substrates (PDL: 1 ± 0, n = 2, N = 3. Flat-ZrOx: 1 ± 0, n = 7, N = 3. ns-ZrOx 1,16 ± 0,16, n = 6, N = 3) (fig. 8e).

**Figure 8:**
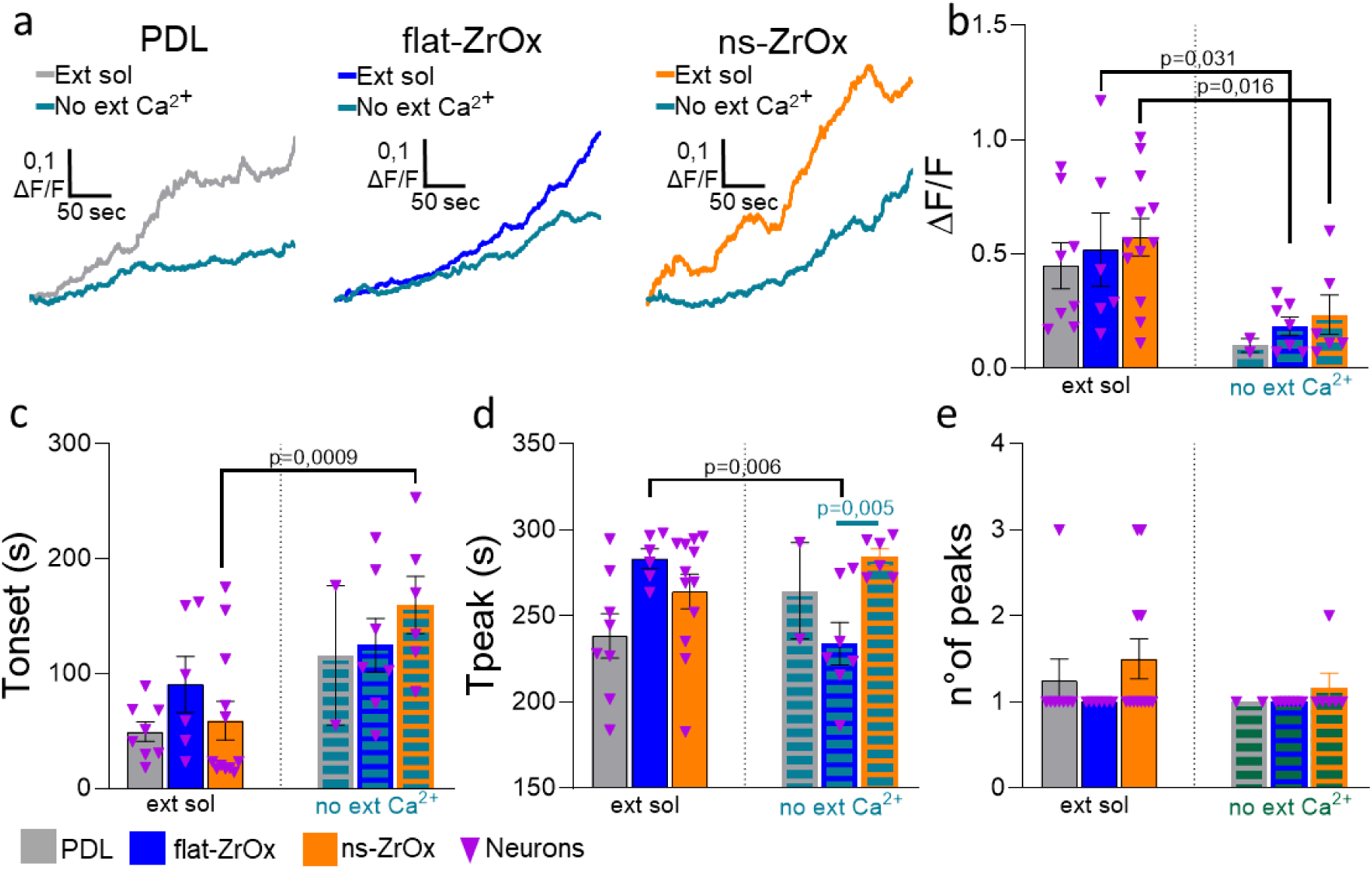
Extracellular calcium contribution on DRG Neuronal calcium responses on ZrOx Substrates. a) Representative traces of fluorescence intensity variation (ΔF/F) over time of DRG neurons plated on all the substrates in control condition and following treatment with 2apb and removal of external Ca²⁺. b) Histogram showing maximum fluorescence intensity values in responding DRG neurons cultured on PDL, flat-ZrOx, and ns-ZrOx in control conditions and removal of external Ca²⁺ [PDL (0,1 ± 0,03, n = 2, N = 3), flat-ZrOx (0,18 ± 0,040, n = 7, N = 3), and ns-ZrOx (0,23 ± 0,08, n = 6, N = 3)]. c) Histogram showing the time onset in responding DRG neurons cultured on PDL, flat-ZrOx, and ns-ZrOx in control conditions and removal of external Ca²⁺ [PDL (115,8 ± 60,75, n = 2, N = 3), Flat-ZrOx (124,8 ± 23,34, n = 7, N = 3), and ns-ZrOx (159,8 ± 24,8, n = 6, N = 3)]. d) Histogram showing the time to peak in responding DRG neurons cultured on PDL, flat-ZrOx, and ns-ZrOx in control conditions and removal of external Ca²⁺ [PDL (264,5 ± 28, n = 2, N = 3), flat-ZrOx (233,9 ± 12,31, n = 7, N = 3), and ns-ZrOx (284,3 ± 4,63, n = 6, N = 3)]. e) Histogram showing the number of peaks in responding DRG neurons cultured on PDL, flat-ZrOx, and ns-ZrOx in control conditions and removal of external Ca²⁺ [PDL (1 ± 0, n = 2, N = 3), flat-ZrOx (1 ± 0, n = 7, N = 3), and ns-ZrOx (1,16 ± 0,16, n = 6, N = 3)]. p values are displayed on the figure.

These data highlight that surface nano-topography not only modulates the amplitude of calcium responses but also their temporal features, in a cell-type-specific manner, reinforcing the potential of ns-ZrOx as a functional interface for glio-neuromorphic applications.

## DISCUSSION

The present study demonstrates that ns-ZrOx films are a biocompatible and functionally active interface for both astrocytes and peripheral neuron-glia co-cultures. By comparing cellular behavior on nanostructured versus flat zirconia substrates, we show that nanoscale topography not only supports cell survival and differentiation but also modulates the amplitude and temporal dynamics of calcium signaling. These effects are consistent across both central (astrocytes) and peripheral (DRG) glial populations, highlighting the potential of ns-ZrOx as a versatile glio-neuromorphic material capable of promoting neuroglial communication.

Astrocytes cultured on ZrOx substrates, both nanostructured and flat, displayed enhanced calcium signaling compared to cells grown on PDL-coated substrates. This enhancement was characterized by statistically significant increased calcium transient amplitude and faster response kinetics. Notably, astrocytes plated on ns-ZrOx exhibited a further potentiation of these effects, indicating that the nanostructured topography amplifies calcium signaling beyond that observed on flat ZrOx surfaces.

Together, these data suggest that ns-ZrOx not only supports astrocyte survival and differentiation but also reshapes their signaling machinery toward a more excitable and responsive state. Such modulation may be advantageous for the design of biomaterials intended to recapitulate the dynamic, bidirectional signaling that characterizes tripartite synapses in vivo.

To extend our study we performed biocompatibility and functional experiments using glial and neuronal cells from the PNS, using a co-culture of DRG neuron and glia which represent a model to study the sensory system. In DRG co-cultures, both neurons and glial cells adhered and survived equally well on all zirconia substrates, confirming the biocompatibility of ZrOx substrates. However, functional imaging revealed that the nanostructured surface selectively enhanced calcium signaling in glial cells, while neuronal activity remained relatively stable across substrates. This selective sensitivity may reflect intrinsic differences in how glia and neurons interact with their microenvironment. Glial cells, being more mechanosensitive and reliant on extracellular cues for calcium wave propagation, likely respond more strongly to the altered topography and surface energy of ns-ZrOx^44,45^. However previous studies showed that when primary hippocampal neurons are cultured on such cluster-assembled nanostructured zirconia surfaces, they display accelerated maturation, earlier action potential generation, and enhanced spontaneous synaptic activity, meaning that ZrOx modulate the functional network formation^46^. Moreover, DRG neurons cultured on nanostructured silicon nanowire mats demonstrated enhanced preservation of functional and electrophysiological activity compared to conventional substrates^47^.

The persistence of partial calcium signaling on ns-ZrOx after IP₃R blockade, in contrast to its complete suppression on flat zirconia and PDL, further highlights the substrate’s influence on intracellular signaling routes. This residual activity may involve alternative calcium sources, such as mechanosensitive channels or store-operated calcium entry (SOCE), whose function could be enhanced by nanoscale features. Similarly, in calcium-free conditions, ns-ZrOx induced distinct temporal changes in both neuronal and glial populations, suggesting that the surface can tune the balance between extracellular influx and intracellular release mechanisms.

These results collectively indicate that ns-ZrOx acts as an active neuromorphic interface, capable of modulating not only neuronal but also glial excitability. The fact that such modulation is consistent across both CNS and PNS cell types underscores the robustness and versatility of zirconia as a glio-compatible material.

Traditionally systems for in materia neuromorphic computing have focused primarily on mimicking neuronal behaviour, replicating spiking, synaptic plasticity. Emerging evidence shows that glial cells play equally critical roles in regulating network excitability and information processing^48–50^. By demonstrating that nanostructured zirconia can differentially affect neuron–glia communication, this study supports the development of glio-neuromorphic interfaces, which integrate both neuronal and glial components to better emulate the complexity of biological information processing.

Moreover, astrocyte-inspired zirconia-based memristive devices have recently been shown to exhibit recoverable conductance linearity and enhanced learning performance, demonstrating how astrocytic principles can be directly translated into hardware implementations to stabilize and improve neuromorphic computation.^51^

The enhanced calcium signaling observed on ns-ZrOx suggests that such substrates can be implemented in hybrid systems that couple living glial–neuronal networks with artificial neuromorphic devices. In particular, the application of a modulated electrical stimuli by the substrates and the recording and processing of cell response by means of the resistive switching activity of the nanostructured material itself can foster neuromorphic bidirectional communication between the artificial and the biological network, in real time and integrated directly in the culture petri. The previously described complex electrical behaviour of ns-ZrOx further reinforce this possibility, indicating that these materials could serve not only as biocompatible scaffolds but also as functional components in signal processing.

More broadly, these findings support an interdisciplinary view of neuromorphic systems in which biological function and device physics must be jointly considered^52^. In this framework, our study provides a direct experimental bridge between glial physiology and nanoscale material properties, positioning nanostructured cluster-assembled thin films as a key enabling platform for next-generation glio-neuromorphic systems.

## CONCLUSIONS

In this work, we identify nanostructured zirconium oxide as a neurogliomorphic interface capable of actively shaping neuroglial function. Beyond its high biocompatibility, ns-ZrOx promotes glial differentiation and selectively enhances glial calcium dynamics, while eliciting distinct responses in neuronal populations. Importantly, our findings show that the effects induced by ns-ZrOx substrate are conserved across both central and peripheral neuroglial systems, indicating that the influence of nanoscale topography on calcium-dependent signaling represents a robust and generalizable phenomenon rather than a specific response. The observation that nanostructured zirconia selectively potentiates glial excitability, while preserving neuronal viability and activity, further supports the concept that glial cells constitute highly responsive targets for neuromorphic material engineering. In this context, ns-ZrOx does not merely function as a passive scaffold, but rather as an active biointerface capable of modulating intracellular signaling pathways and neuroglial communication. These features are particularly relevant for the development of hybrid bioelectronic systems, where the bidirectional interaction between living neural networks and functional materials requires dynamic and adaptive signaling capabilities.

Overall, this work contributes to a broader shift from neuron-centered approaches toward integrated neuron–glia paradigms that more faithfully reproduce the emergent properties of biological neural networks. By bridging glial physiology and neuromorphic nanostructured materials, our study establishes cluster-assembled zirconia thin films as a promising platform for next-generation biohybrid technologies, including adaptive neural interfaces, brain-inspired computing architectures, and real-time neuroelectronic communication systems.

## AUTHOR INFORMATION

### Author Contributions

The manuscript was written through contributions of all authors. All authors have given approval to the final version of the manuscript. V.B., P.M., G.C. and F.B. conceived and designed the experiments.C.P. fabricated flat- and ns-ZrOx substrates. F.B. characterised the substrates, G.C. and C.L. prepared the cellular cultures, performed immunofluorescence characterization, viability test, calcium imaging and analyzed the related data. G.C., C.L., A.K. and R.F. prepared and maintained the astrocytes and DRG cell culture. M.C. supported the preparation of the cell cultures. G.C wrote the manuscript. All authors discussed the results and contributed to the manuscript writing and reviewing.

## ABBREVIATIONS

ZrOx: zirconium oxide
ns: nanostructured
DRG: dorsal root ganglion
([Ca^2+^]_i_): intracellular calcium signaling
ECM: extracellular matrix
ns-ZrOx/Au: nanostructured zirconia and gold
SCBD: Supersonic Cluster Beam Deposition
PMCS: Pulsed Micro Plasma Cluster Source
Rq: roughness
AFM: Atomic Force Microscopyy
DMEM: Dulbecco’s Modified Eagle Medium
FBS: fetal bovine serum
FDA: Fluorescent Diacetate Assay
PDL: Poly-D-lysine
DIV: days in vitro
2APB: 2-Aminoethyl Diphenylborinate
IP_3_: inositol 1,4,5-trisphosphate
(0Ca^²⁺^): calcium-free extracellular saline
CNS: central nervous system
PNS: peripheral nervous system
GAP-43: Growth Associated Protein-43
SOCE: store-operated calcium entry.

